# The landscape of alternative polyadenylation during EMT and its regulation by the RNA-binding protein Quaking

**DOI:** 10.1101/2022.12.01.518773

**Authors:** Daniel P. Neumann, Katherine A. Pillman, B. Kate Dredge, Andrew G. Bert, Cameron P. Bracken, Brett G. Hollier, Luke A. Selth, Traude H. Beilharz, Gregory J. Goodall, Philip A. Gregory

**Affiliations:** Centre for Cancer Biology, University of South Australia and SA Pathology, Adelaide, SA, 5000, Australia; Faculty of Health and Medical Sciences, The University of Adelaide, Adelaide, SA 5000, Australia; Australian Prostate Cancer Research Centre - Queensland, Centre for Genomics and Personalised Health, Faculty of Health, School of Biomedical Sciences, Queensland University of Technology, Princess Alexandra Hospital, Translational Research Institute, 37 Kent Street, Woolloongabba, Brisbane, QLD, 4102; Flinders Health and Medical Research Institute, Flinders University, Bedford Park, SA 5042, Australia; Development and Stem Cells Program, Monash Biomedicine Discovery Institute and Department of Biochemistry and Molecular Biology, Monash University, Melbourne, VIC 3800, Australia

**Keywords:** Quaking, epithelial-mesenchymal transition, alternative polyadenylation

## Abstract

Epithelial-mesenchymal transition (EMT) plays important roles in tumour progression and is orchestrated by dynamic changes in gene expression. While it is well established that post-transcriptional regulation plays a significant role in EMT, the extent of alternative polyadenylation (APA) during EMT has not yet been explored. Using 3’ end anchored RNA sequencing, we mapped the alternative polyadenylation landscape (APA) following TGF-β-mediated induction of EMT in human mammary epithelial cells and found APA generally causes 3’UTR lengthening during this cell state transition. Analysis of the RNA-binding protein Quaking (QKI), a splicing factor induced during EMT, revealed enrichment of its binding adjacent to cleavage and polyadenylation sites within 3’UTRs. Following QKI knockdown, APA of many transcripts are altered to produce predominantly shorter 3’UTRs associated with reduced gene expression. Among these, QKI binds to its own cleavage site to produce a transcript with a longer 3’UTR. These findings reveal extensive changes in APA occur during EMT and identify a novel function for QKI in this process.

## Introduction

Epithelial-mesenchymal transition (EMT) is a developmental process that facilitates cancer progression in epithelial-derived tumours, through the acquisition of mesenchymal features (Yang et al. 2020). EMT can be initiated by extracellular cues that regulate extensive changes in gene transcription through induction of EMT transcription factors (De Craene and Berx 2013). Of note, TGF-β signalling is a potent inducer of the transcription factors ZEB1 and ZEB2, which operate in a reciprocal feedback loop with miR-200 to drive dynamic changes in gene expression and cell state (Bracken et al. 2008; Burk et al. 2008; Gregory et al. 2008; Gregory et al. 2011). EMT also involves widespread changes in alternative splicing mediated by specific RNA-binding proteins (RBP) (Shapiro et al. 2011; Yang et al. 2016; Harvey et al. 2018; Neumann et al. 2018). Recently, we and others identified a prominent role for the RBP Quaking (QKI) in promoting EMT-associated alternative splicing and the acquisition of mesenchymal cell traits through maintenance of mesenchymal splicing patterns (Li et al. 2018; Pillman et al. 2018). QKI expression is elevated by loss of miR-200 during TGF-β driven EMT, where it binds to introns and results in the alternative splicing of hundreds of mRNA targets (Pillman et al. 2018).

QKI is itself alternatively spliced to produce three major isoforms, QKI-5, QKI-6 and QKI-7, with distinct functions. By virtue of a nuclear localisation signal, QKI-5 is localised in the nucleus, while QKI-6 and QKI-7 are predominantly expressed in the cytoplasm (Wu et al. 1999). QKI isoforms regulate many aspects of mRNA metabolism in addition to alternative splicing including mRNA stability, localisation and translation (Larocque et al. 2002; Larocque et al. 2005; Zhao et al. 2006; Yamagishi et al. 2016; Sakers et al. 2021; Neumann et al. 2022). Consistent with these functions, a subset of QKI-binding occurs within 3’UTRs during EMT (Pillman et al. 2018), however, it is currently unclear what the functional consequences of QKI 3’UTR binding are in mesenchymal cells.

The influence of alternative cleavage and polyadenylation (APA) on transcript diversity during EMT is largely unexplored. APA can produce transcripts with varying 3’UTRs or alterative last exons, which can impact mRNA stability, localisation, translation and in some cases regional translation of specific mRNAs (Mitschka and Mayr 2022). RBPs can co-transcriptionally regulate APA leading to coordination of APA in a cell specific manner based on the relative expression or activities of these factors (Tian and Manley 2017). While 3’UTR shortening is prevalent in solid cancers and can promote expression of oncogenes (Mayr and Bartel 2009; Xia et al. 2014), broader comparisons of APA and gene expression changes show no consistent relationship, indicating that these two mechanisms of gene regulation operate largely independently (Lianoglou et al. 2013; Wang et al. 2018).

Here, we performed 3’ end anchored RNA sequencing (PolyA-Test RNA-seq or PAT-Seq) to quantitate the landscape of APA during EMT. Following TGF-β-induced EMT of human mammary epithelial cells (HMLE), we identified 79 events with significant APA, two-thirds of which resulted in longer 3’UTRs. By analysis of QKI CLIP-seq data, we identified a large proportion of QKI 3’UTR binding occurs adjacent to productive cleavage and polyadenylation (CPA) sites and found QKI knockdown in mesenchymal HMLE (mesHMLE) cells generally shortens 3’UTRs. These findings reveal the APA landscape during EMT and highlight a novel function for QKI as a regulator of APA.

## Results

### Alternative polyadenylation events regulated during EMT and by loss of QKI

While it is well established that post-transcriptional regulation plays a significant role in EMT, the extent of APA during EMT has not yet been explored. As the identification of shifts in CPA site usage requires a 3’end anchored RNA-Seq method, we performed PAT-Seq (Harrison et al. 2015; Swaminathan et al. 2019) on triplicate samples of human mammary epithelial (HMLE) cells and HMLE cells treated with TGF-β for 20 days to induce EMT (mesHMLE cells). We also performed PAT-seq on mesHMLE cells transfected with siRNAs targeting QKI-5, a known regulator of alternative splicing in EMT, but with no recognised role in APA (Pillman et al. 2018), generating on average ~8 million mapped reads per sample (Supplementary Table 1). Multidimensional scaling analysis showed distinct clustering of each sample type, with the exception of one mesHMLE QKI siRNA sample (R3 – transfected with a different QKI siRNAs to R1 and R2), which was excluded from further analysis (Supplementary Figure 1A & Supplementary Table 2). This technique enabled us to identify changes in CPA site usage and poly(A) tail length during EMT as well as those regulated by QKI.

We verified the induction of EMT in the mesHMLE samples by confirming downregulation of epithelial (E-cadherin, ESRP1) and upregulation of mesenchymal (ZEB1, Vimentin, N-cadherin, and QKI) markers (Supplementary Figure 1B, Supplementary Table 3 & 4). Transfection of mesHMLE with the QKI siRNA led to reduced QKI expression and did not change EMT markers, as previously observed (Supplementary Figure 1B) (Pillman et al. 2018). In each group of samples, the mean poly(A) tail length was similar indicating global changes are not a feature of EMT or regulated by QKI (Supplementary Table 1). Likewise, tests for changes in polyA tail length of specific genes, during EMT or after QKI knockdown did not uncover any significant events (Supplementary Table 5 & 6). In contrast, there were 79 genes with significantly affected APA during EMT, with 27 shortening events and 52 lengthening events (Figure 1A, Supplementary Table 7). QKI knockdown resulted in an opposing pattern of APA with 94 shortening and 25 lengthening events (total 119 genes, Figure 1B, Supplementary Table 8), suggesting QKI may promote a shift to distal CPA site usage during EMT. Examining the intersection in these datasets showed little overlap between the statistically significant APA events (7 genes), although each of these events changed in opposing directions (Supplementary Figure 1C). This data suggests that QKI regulates a subset of APA events during EMT, potentially similar to its role in regulating EMT associated alternative splicing (Pillman et al. 2018).

**Figure 1:**
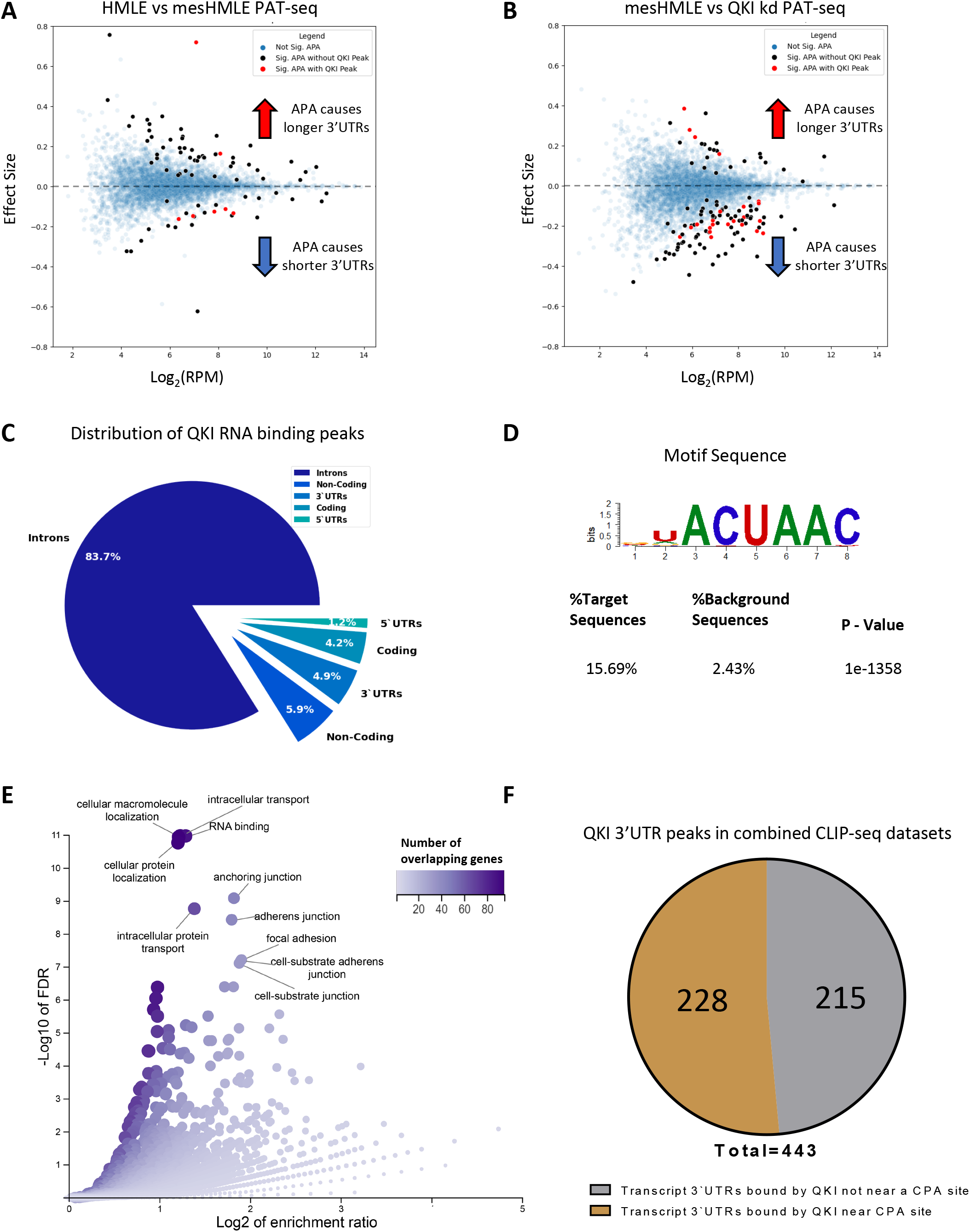
PAT-Seq uncovers widespread switching to distal cleavage sites after EMT and proximal sites after QKI knockdown. (A) MA plots for HMLE vs mesHMLE with APA effect size plotted against average expression across samples. (B) MA plot for mesHMLE vs mesHMLE + QKI knockdown. A positive effect size value indicates a shift to distal cleavage site usage, while a negative value indicates a shift to proximal sites. Blue dots indicate no significant change in APA, black dots indicate a significant change in APA and red dots indicate a significant change in APA and a QKI CLIP-Seq peak present in the 3’UTR of that transcript (in at least 3 QKI CLIP-Seq replicates). (C) Pie chart showing the distribution of QKI CLIP-Seq peaks by genomic location type. (D) The most enriched motif in the mesHMLE HITS-CLIP as determined by the software package HOMER. (E) Volcano plot of enriched GO terms from the set of 443 transcripts bound by QKI in at least 4 out of 7 separate CLIP-Seq experiments. Dot colour and size represents number of overlapping genes for that category. (F) Pie chart depicting the proportion of QKI 3’UTR targets bound within 100 nucleotides of an annotated cleavage site from the Poly A site atlas.

### QKI binds near productive cleavage and polyadenylation sites

Previously, we uncovered numerous QKI binding sites within introns by performing QKI-5 HITS-CLIP on mesHMLE cells (Pillman et al. 2018). These sites reflect QKI’s established role as a regulator of alternative splicing, where QKI-5 was found to directly bind and regulate alternative splicing of hundreds of mRNAs during EMT. However, a subset of peaks were found in 3’UTRs, suggesting that QKI may regulate some mRNAs through non-splicing mechanisms (Figure 1C).

CLIP-Seq results can be greatly affected by the choice of methods and antibodies, and the choice of peak calling requires the selection of parameters that strike a balance between sensitivity and stringency (Hafner et al. 2021). Therefore, we sought to generate a list of high confidence QKI 3’UTR targets by pooling data from several QKI CLIP-Seq experiments. We performed QKI HITS-CLIP using two separate QKI antibodies, one specific for QKI-5 and another that recognized an epitope common to all QKI isoforms (panQKI). Consistent with our previous observations (Pillman et al. 2018), we identified the known QKI motif to be the most highly enriched sequence within the peaks (Figure 1D) and found the majority of QKI binding occurred within introns (Figure 1C).

Within 3’UTRs, we identified 429 distinct mRNAs with peaks suggesting that QKI could regulate these transcripts (Supplementary Table 9). However, only 157 of the 3’UTRs with peaks in the panQKI HITS-CLIP experiment had peaks in our previously published QKI-5 HITS-CLIP data (Supplementary Figure 2A). This could be indicative of false positive or negative peaks or cytoplasmic-specific QKI interactions since the panQKI antibody recognises both cytoplasmic and nuclear isoforms of QKI. To generate a list of putative QKI 3’UTR targets with a greater level of confidence, we expanded our analysis to include various published QKI CLIP-seq datasets. We analysed data from PAR-CLIP performed on HEK293 human embryonic kidney cells (PAR-CLIP-HEK293) and eCLIP performed on K562 chronic myelogenous leukemia cells (K562-rep-1 and K562-rep-2) and HepG2 hepatocellular carcinoma cells (HEPG2-rep-1 and HEPG2-rep-2) (Hafner et al. 2010; Van Nostrand et al. 2016), constraining our analysis to peaks called within 3’UTRs (Supplementary Figure 2B, 2C). Using these datasets, we generated a unique list of regions that overlap with a peak in at least one QKI CLIP-Seq experiment (Supplementary Table 10). We subsequently tallied the number of experiments where a peak was successfully called for each unique region, and shortlisted regions containing a peak in at least four out of the seven experiments. This resulted in the identification of 622 peak-containing regions that were present within 443 distinct transcripts (Supplementary Table 10). To determine if these genes have functions that are relevant to epithelial plasticity, we performed gene ontology (GO) analysis and uncovered pathways related to EMT-associated properties, including cell junctions and focal adhesions (Figure 1E), supporting the notion that regulation of these mRNAs by QKI might be functionally important.

Next, we explored whether QKI binding within 3’UTRs may be involved in regulating APA. Recent analysis of the cytoplasmic isoform, QKI-6, found an enrichment of binding within 3’UTRs adjacent to the polyA signal (PAS) (Sakers et al. 2021). As the QKI isoforms share the same RNA-binding domain and recognize the same RNA-binding motif (Galarneau and Richard 2005), we speculated that the nuclear isoform, QKI-5, might also bind near PAS and affect CPA. To assess this possibility, we compared the QKI CLIP-Seq data to the PolyA site atlas (Herrmann et al. 2020) to identify peaks that occur near productive CPA clusters. Of the 433 QKI 3’UTR targets, approximately half had annotated CPA sites within or nearby a QKI CLIP-Seq peak (Figure 1F). These results suggest that a large proportion of QKI 3’UTR-binding occurs at productive CPA sites and uncovers a putative role for QKI in regulating alternative polyadenylation.

### A subset of alternative polyadenylation during EMT is controlled by QKI

To uncover APA events directly regulated by QKI, we compared the genes with altered APA after QKI knockdown to QKI 3’UTR CLIP-seq targets, identifying 24 genes that had altered APA and evidence of QKI binding in at least 3 separate experiments. This overlap was highly significant (P= 1.05 × 10^−9^, hypergeometric test), suggesting QKI directly regulates APA of this subset of mRNAs (Figure 2A). Gene ontology (GO) of these genes highlighted processes including lamellipodium, fibroblast migration and cell leading edge, indicating that QKI-regulated APA may contribute to changes in cell migration (Figure 2B).

**Figure 2:**
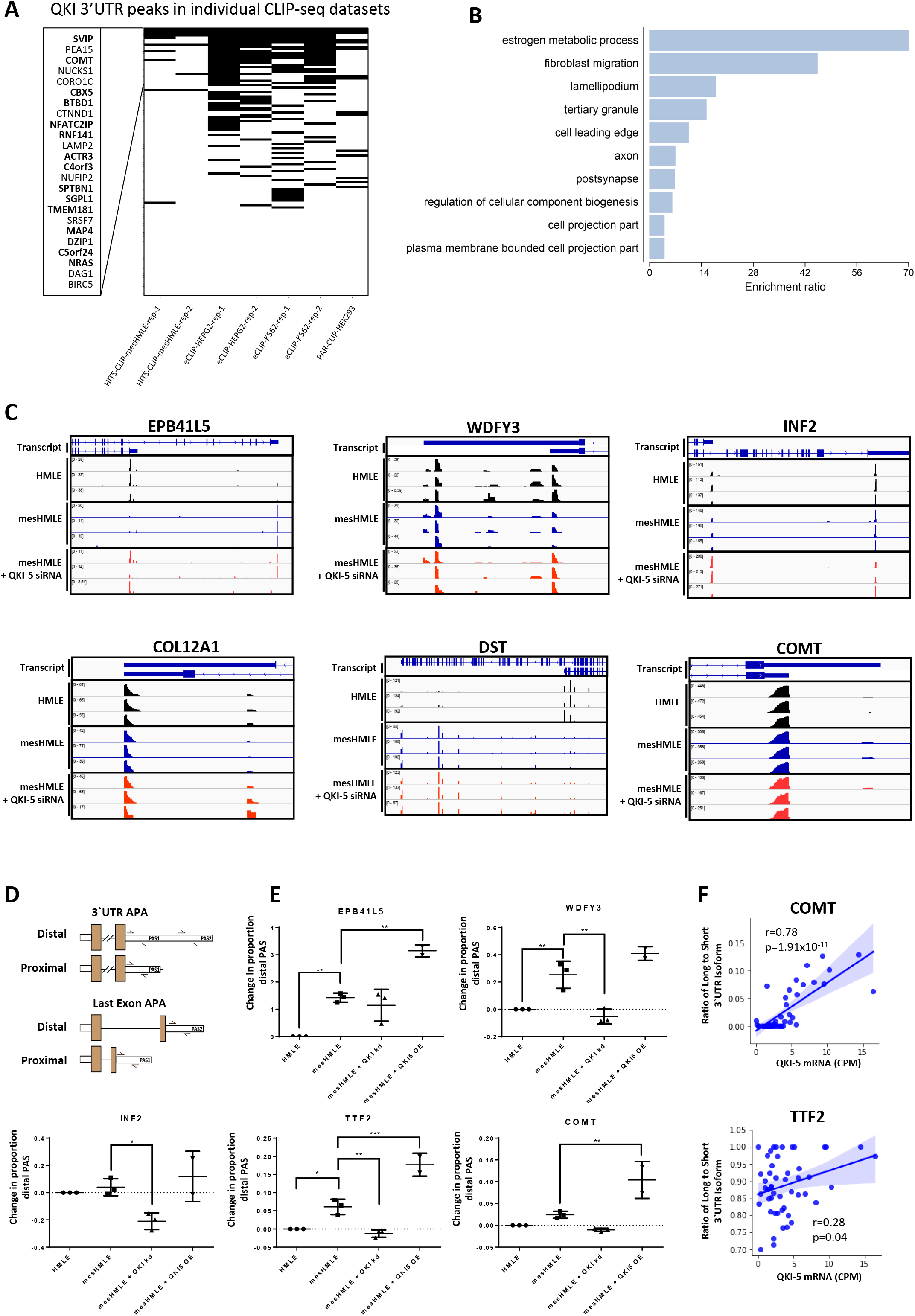
QKI 3’UTR binding often occurs near cleavage and polyadenylation sites. (A) Heat map comparison of QKI regulated APA events with QKI CLIP-seq datasets to identify APA events directly regulated by QKI. Genes with APA events (n=24) and evidence of QKI binding in at least 3 QKI CLIP-seq datasets are highlighted in bold. (B) Bar chart of enriched GO terms in the set of 24 genes. (C) Images of IGV tracks depicting change in cleavage site usage for selected genes. Individual tracks show depth-normalized coverage of each replicate PAT-Seq sample for HMLE, mesHMLE and mesHMLE + QKI knockdown. (D) Graphs showing the change in APA site usage for selected genes in HMLE (n=3), mesHMLE (n=3), mesHMLE + QKI-5 knockdown (n=3) and mesHMLE + QKI-5 overexpression (n=2), as determined by qRT-PCRs. A schematic of primer locations is depicted, with plots showing the change in proportion of distal cleavage to proximal cleavage. For COMT, INF2, TTF2, WDFY3 and EPB41L5, proximal cleavage primers detect proximal and distal APA and are a measure of total mRNA. Error bars indicate standard deviation of the mean. Statistical significance determined by performing a one-way ANOVA (p<0.05), followed by Sidak’s multiple comparison post-hoc test. *p<0.05, **p<0.01, ***p<0.001. (E) Scatter plot showing the ratio of the Long to short 3’UTR transcripts against total QKI mRNA in the Cancer Cell Line Encyclopedia breast cancer samples (Barretina et al. 2012). Line represents a linear regression model and the shaded area represents the confidence interval. Pearson correlation coefficients and associated p-values for COMT and TTF2 are shown.

To validate the findings of the PAT-Seq, we performed qRT-PCR on a set of candidate genes with significantly altered APA during EMT or following QKI knockdown (Figure 2C). We also included genes that, although not significantly altered with QKI knockdown in the PAT-seq, had strong evidence of QKI binding near to a productive CPA site. To detect a switch between mRNA isoforms with different cleavage sites within the same 3’UTR, we designed a primer set specific for the longer 3’UTR and another set that amplifies a common region. For cleavage sites in distinct 3’UTRs, we designed primers that were specific for each isoform (Figure 2D). We found several genes with consistently altered APA after EMT, that changed in the opposite direction after QKI knockdown or were augmented after QKI-5 overexpression in mesHMLE cells, consistent with QKI controlling the choice between the alternative CPA sites (Figure 2E).

To determine whether the effects of QKI on APA occur more broadly, we examined expression of long and short 3’UTR isoforms in breast cancer cell lines within the Cancer Cell Line Encyclopaedia (Barretina et al. 2012). We found a positive correlation between the ratio of long to short 3’UTR transcript and *OKI-5* mRNA for *COMT* and *TTF2* suggesting that the balance between the long and 3’UTRs of these genes is dependent on QKI expression (Figure 2F).

### QKI regulates alternative polyadenylation of its own transcript

Among the QKI-regulated APA genes, we identified two CLIP-seq peaks within the *OKI* transcript itself that overlapped canonical QKI-binding motifs and were located immediately downstream from an annotated CPA site for QKI-5 (Figure 3A). This region encompasses three distinct polyadenylation signals (PAS), one of which appears to be the predominant signal for polyadenylation of QKI-5 (Figure 3B). When examining the *OKI-5* 3’UTR in the UCSC genome browser, we found that many expressed sequence tags (ESTs) ended at the short 3’UTR terminus (QKI-5-Short) identified previously (Pillman et al. 2018) or ~2kb upstream, with a smaller number ending at the annotated long 3’UTR terminus (QKI-5-Long) (Supplementary Figure 3A). At the QKI-5-Short terminus, the majority of ESTs ended at the QKI binding site downstream of the predominant PAS (Supplementary Figure 3B), which lead us to hypothesize that QKI may repress cleavage and polyadenylation at this location and utilise a downstream PAS to generate a longer QKI-5 3’UTR.

**Figure 1:**
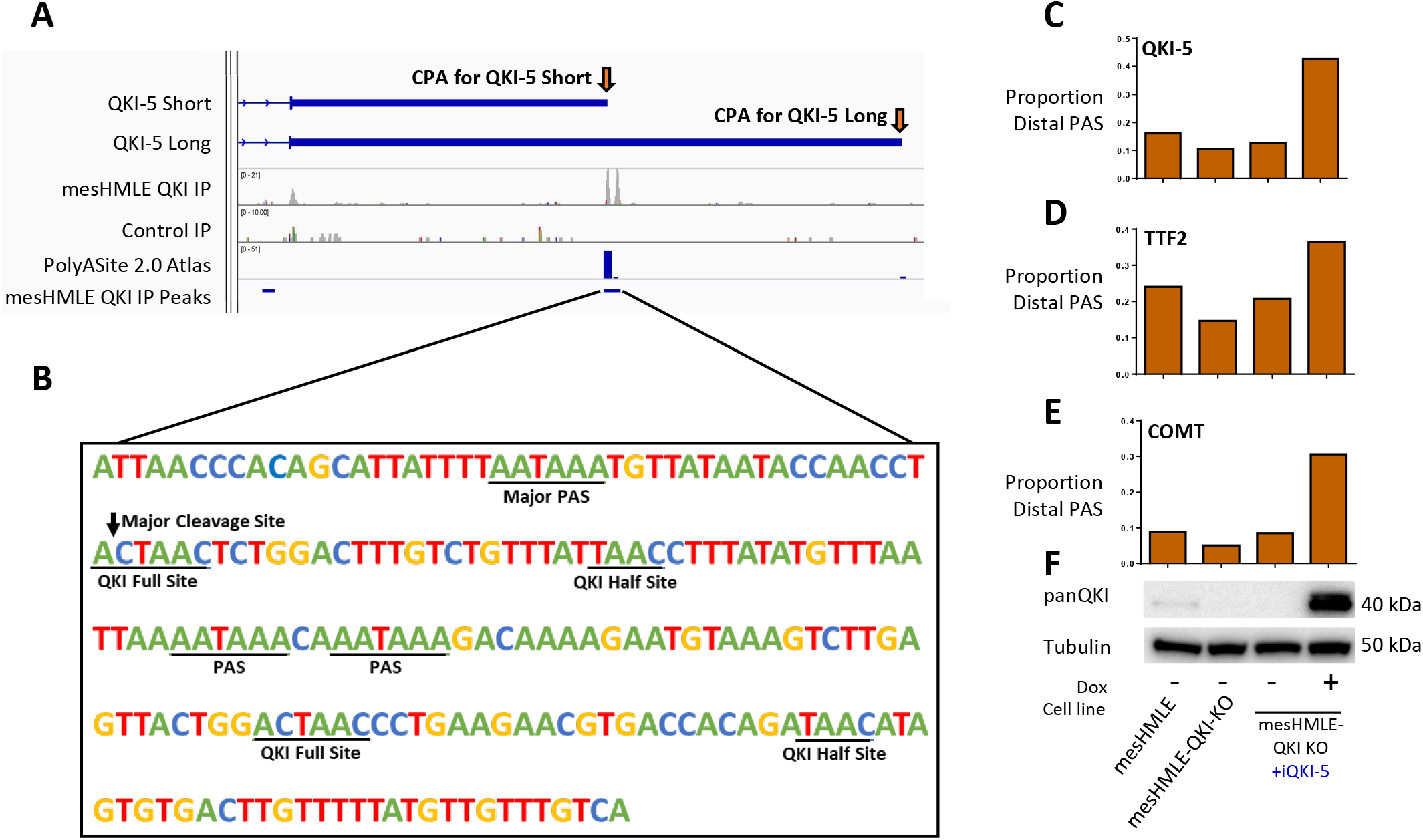
QKI-5 regulates its own APA to produce a transcript with a long 3’UTR. (A) IGV tracks depicting the location of QKI CLIP-Seq peaks in the QKI-5 3’UTR. (B) The nucleotide sequence around the predominant site of QKI binding in the QKI-5 3’UTR showing potential QKI binding and PAS sites. (C-E) Bar charts depicting the ratio of the isoform that uses the distal cleavage site to the isoform that uses the proximal cleavage sites for indicated genes determined by qRT-PCRs performed on mesHMLE, mesHMLE QKI knockout and mesHMLE QKI Knockout + inducible QKI-5, with and without doxycycline treatment. (F) Western blots of QKI and Tubulin protein performed on matched protein samples for (C-E).

To confirm the existence of a *OKI-5* transcript with a longer 3’UTR and determine if QKI-5 protein induces its production, we examined expression of the QKI-5-Short and QKI-5-Long in a mesHMLE clonal cell line with CRISPR-generated QKI knockout. These cells had reduced QKI protein expression but retained QKI-5 mRNA, allowing us to modulate QKI-5 protein levels by exogenous expression and measure effects on endogenous QKI-5 mRNA. We transduced this cell line with a doxycycline-inducible QKI-5 expression construct and measured QKI-5-Long and total QKI in wild-type mesHMLE, QKI knockout, and QKI knockout cells with or without QKI-5 overexpression. Reduced QKI-5 protein in the knockout cells led to decreased production of QKI-5-Long mRNA, but not total endogenous QKI-5 mRNA, and a consequent decrease in distal to proximal PAS usage (Supplementary Figure 4, Figure 3C and 3F). In contrast, overexpression of QKI-5 increased QKI-5-Long while reducing total endogenous QKI-5 mRNA, leading to an increase in the distal to proximal PAS usage (Supplementary Figure 4, Figure 3C and 3F). In these samples, we also verified the proportion of distal to proximal PAS usage for *COMT* and *TTF2* are also influenced by QKI levels as observed previously (Figure 3D and 3E). Collectively, these findings indicate that QKI-5 regulates its own APA, and the APA of other genes, to generate isoforms with generally longer 3’UTRs during EMT.

### QKI regulated APA increases 3’UTR length and gene expression

To investigate the potential consequences of APA during EMT, we examined the relationship of APA to gene expression changes. While EMT was associated with large changes in gene expression (2203 genes, FDR <0.05, Supplementary Figure 5A and Supplementary Table 3), only modest numbers of genes change expression following QKI knockdown (101 genes, FDR <0.05, Supplementary Figure 5B and Supplementary Table 4), consistent with previous observations (Pillman et al. 2018). Of the 79 genes with changes in APA during EMT, 38 also had significant changes in gene expression, however there was no consistent relationship between the direction of change with either 3’UTR shortening or lengthening (Figure 4A and 4C). However, 28 of the 119 QKI-regulated APA events had significant changes in gene expression, with 23 of these having shorter 3’UTRs and 19 of these being downregulated (Figure 4B and 4D). Collectively, these data suggest that QKI-regulated APA generally increases 3’UTR length and gene expression in a subset of genes during EMT.

**Figure 4:**
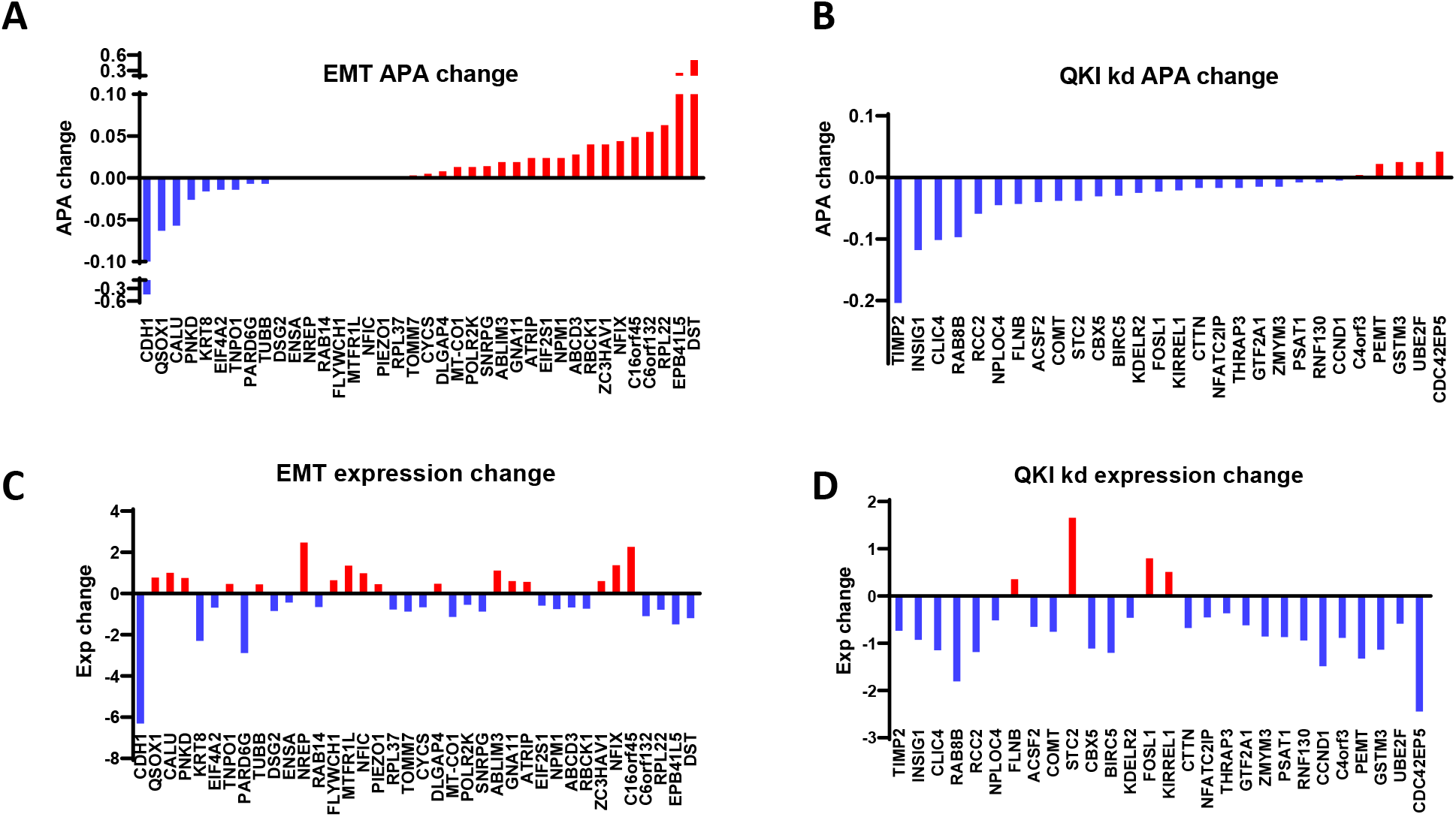
Correlation of APA changes with gene expression changed during EMT and following QKI knockdown. (A-B) Changes in APA during EMT of HMLE cells or following QKI knockdown in mesHMLE cells, aligned from most shortened to most lengthened 3’UTR. Only genes with significant changes in APA (FDR<0.05) and gene expression (p<0.05) are represented (C-D) Corresponding changes in gene expression for APA affected genes. Blue bars indicated decreased APA and gene expression, and red bars indicate increased APA and gene expression.

## Discussion

To our knowledge, this study represents the first targeted examination of alternative polyadenylation (APA) during EMT using 3’ end anchored RNA sequencing. Here, we have shown that many transcripts display APA after EMT in human mammary epithelial cells, and present evidence that the mesenchymal splicing factor QKI regulates APA of several transcripts during EMT, including its own, to produce mRNAs with distinct 3’UTRs or last exons. This supports the existence of a mesenchymal-specific APA signature that is invoked by QKI and warrants further investigation of how widespread this mode of regulation is in other contexts of EMT and cellular differentiation regulated by QKI (Neumann et al. 2022).

With the PAT-seq method, we detected many APA events (~200) that were regulated during EMT and/or directly by QKI. While there was a small overlap between EMT and QKI knockdown regulated APA events, this is not entirely unexpected given that APA can be influenced by many RBPs, including those that are regulated during EMT (Shapiro et al. 2011; Dittmar et al. 2012). However, validation of APA events by qPCR indicates that a larger fraction may be regulated in opposing directions in both scenarios. For instance, while the ratio of the Long to Short 3’UTRs of COMT and TTF2 increased during EMT and responded to QKI manipulation, these ratios were not significantly changed in the EMT PAT-seq dataset. This highlights limitations of the sensitivity of PAT-seq at the sequencing depth reported here (5-10 million reads per sample mapping to the genome) and suggests that the spectrum of QKI regulated APA events during EMT may be even larger.

While many APA events were detected to be regulated by QKI, we constrained our analysis to events where direct evidence for QKI 3’UTR binding was observed. Gene Ontology of this select gene subset revealed enrichment of EMT-associated properties indicating APA may cause functional changes consequential for EMT. For example, both INF2 and EBP41L5 have reported functions in controlling focal adhesion and actin dynamics that are important for cell migration (Chhabra and Higgs 2006; Schell et al. 2017). Discovery of how these alternative 3’UTR sequences affect gene expression during EMT warrants further study.

During EMT, approximately two-thirds of APA events were associated with 3’UTR lengthening, while the opposite pattern occurred upon QKI knockdown in mesenchymal cells. Examination of the mRNA levels of APA regulated transcripts during EMT revealed no correlation between gene expression and APA. In contrast, QKI knockdown generally decreased the expression of mRNAs whose APA it directly affected. Given QKI knockdown only influenced overall gene expression modestly, this suggests QKI regulated APA may play a prominent role in generating more stable mRNA transcripts. Alternatively, as the binding of cytoplasmic forms of QKI are known regulate the stability of specific transcripts (Larocque et al. 2005; Zhao et al. 2006; Doukhanine et al. 2010; Thangaraj et al. 2017; Sakers et al. 2021), QKI may influence both APA and mRNA stability through different mechanisms. Although APA induced 3’UTR shortening is a feature of cancer cells and can induce oncogene expression (Mayr and Bartel 2009), broad associations between APA and gene expression are generally not observed (Lianoglou et al. 2013; Brumbaugh et al. 2018; Wang et al. 2018). APA and associated changes in 3’UTR usage can alter the localization and local translation of mRNA transcripts (Lau et al. 2010). Considering the dynamic nature of epithelial plasticity, and the observation that altered protein localization can drive features of EMT in cancer (Aiello et al. 2018), it may be that changes in APA of these mRNAs change the fate of their coded proteins, with potential consequences for protein folding, complex formation and localization. Indeed, protein complex formation and localization have been suggested to be 3’UTR-directed, with the long 3’UTR variant of CD47, specifically, interacting with SET and localizing to the plasma membrane (Berkovits and Mayr 2015). Further research is needed to determine what the functional consequences of QKI-directed APA are to the target mRNAs coded protein.

While the finding that QKI regulates its own APA has not been previously reported, the ability of QKI to self-regulate is known (Darbelli et al. 2017; Fagg et al. 2017). Indeed, the tendency of RBPs to auto-regulate is a well-established phenomenon, and auto-regulation of APA has been reported for other proteins of this class (Muller-McNicoll et al. 2019). Whether QKI alternative polyadenylation serves as a mechanism of negative feedback or the switch to a longer 3’UTR affects the localization or function of QKI protein will require further investigation.

In summary, we have uncovered a novel role for QKI in regulating extensive APA of transcripts during EMT. Along with its function in regulating EMT associated alternative splicing (Pillman et al. 2018), these data demonstrate the diverse roles of QKI in regulating transcript maturation during EMT.

## Acknowledgements

We acknowledge the assistance of Angavai Swaminathan for PAT-seq library preparation, and Paul Harrison and the Monash Bioinformatics Platform for PAT-seq data management. This work was supported by grants from the National Health and Medical Research Council of Australia to P.A.G. and G.J.G. (1128479, 1164669), and the Hospital Research Foundation. T.H.B. is supported by the ARC (FT180100049). G.J.G is supported by a National Health and Medical Research Council fellowship (1118170). P.A.G. and L.A.S. are supported by Principal Cancer Research Fellowships awarded by Cancer Council’s Beat Cancer project on behalf of its donors, the state Government through the Department of Health, and the Australian Government through the Medical Research Future Fund.

## Materials and Methods

### Cell culture

The immortalized human mammary epithelial cell line (HMLE) was cultured in HuMEC Ready Media (ThermoFisher) with HuMEC Supplement Kit (ThermoFisher) and passaged at a 1:5 ratio every 72 hours. The TGF-β-treated mammary epithelial cell line (mesHMLE) was derived by culturing immortalized human mammary epithelial cells (HMLE) in mesMHLE culturing media (DMEM/F12-Dulbecco’s Modified Eagle Medium: Nutrient Mixture F-12 supplemented with 5% Foetal Bovine Serum, 20 ng/ml EGF, 10 μg/ml insulin and 0.5 μg/ml hydrocortisone) with 2.5 ng/μL TGF-β1 for at least 14 days (Mani et al. 2008). Once established, mesHMLE and mesHMLE QKI KO cells were all cultured in mesHMLE culturing media and passaged at a 1:10 ratio every 72 hours.

### Generation of QKI knockout and overexpression cell lines

QKI knockout cells were generated by CRISPR-Cas9 gene editing. A guide RNA sequence designed to target the first exon of the QKI gene (Supplementary Table 11) was combined with tracrRNA and Cas9 protein (Integrated DNA Technologies, AltR system) according to the manufacturer’s protocol and transfected into mesHMLE using Lipofectamine RNAiMAX (ThermoFisher). Single clones were isolated following 72 hr of culture and screened for loss of QKI protein by Western Blot. QKI-5 overexpression cells were generated by transduction of mesHMLE wild-type or QKI ko cells with a plnducer20-QKI-5 lentivirus generated as previously described (Pillman et al. 2018).

### Transfection of siRNAs

Transfections in this study were carried out with 20 nM of siRNA using Lipofectamine RNAiMAX and performed according to the manufacturer’s protocol (ThermoFisher). Transfection media was removed the following day after transfection and cellular material was harvested 72 hours after transfection. Sequences of siRNAs are provided in Supplementary Table 12.

### RNA Isolation, cDNA synthesis and qRT-PCR

RNA was extracted using Trizol (ThermoFisher) following the standard manufacturer’s protocol. Complementary DNA was synthesized from 1 μg of RNA using the QuantitTect RT kit (Qiagen). Quantitative PCRs (qPCR) were performed in triplicate using the QuantiTect SYBR Green reagent (Qiagen) on a Rotor-Gene 6000 series thermocycler (Qiagen). Analysis was performed using the comparative quantitation feature in the Rotor-Gene software and qPCR assays were normalized to GAPDH expression to determine relative mRNA expression. To determine the proportion of distal polyadenylation site (PAS) usage for a given gene, the expression of the distal isoform is divided by the total expression of the gene. Sequences for the oligonucleotides used in the study are listed in Supplementary Table 11.

### Protein lysate purification and western blotting

Protein extracts were obtained by lysis of cells in 1xRIPA buffer (Abcam) containing complete Mini, EDTA-free Protease inhibitor Cocktail tablets (Roche) and phosphatase (PhosSTOP EASYpack (Roche) inhibitors. Twenty micrograms of lysate were separated on an Invitrogen Bolt Bis-Tris Plus gel (Life Technologies), transferred to nitrocellulose, and probed with α-Tubulin (Abcam, ab729l) and ct-panQKI (Neuromab, N147/6) antibodies diluted 1:5000 in 5% skim milk. Proteins were detected by enhanced chemiluminescence (ECL) reagent (Pierce) using the ChemiDoc imaging system (Bio-Rad).

### QKI HITS-CLIP

QKI HITS-CLIP was performed as detailed previously (Pillman et al. 2018). QKI was immunoprecipitated with a QKI-5 specific (Bethyl, A300-183A) or panQKI (Neuromab, N147/6) antibodies and sequencing libraries prepared including a size matched input control (Van Nostrand et al. 2016).

### PolyA-Test RNA-seq (PAT-seq)

PAT-seq libraries were prepared using 1 ug of total RNA as previously described (Harrison et al. 2015; Swaminathan et al. 2019). Libraries were sequenced multiplexed on the Illumina NovaSeq 6000 instrument (Alfred Research Alliance Genomics Facility). Raw data was processed using the tail-tools pipeline (Harrison et al. 2015). The data from all samples was combined to identify 25437 peaks associated with non-templated poly A-tracts. Of these, 20928 peaks and 73.3% of reads were in a 3’ UTR, 1349 peaks and 16.1% of reads were otherwise in an exon, 1202 peaks and 3.8% of reads were downstream of a non-coding RNA, 957 peaks and 0.7% of reads were in an intron, 403 peaks and 0.2% of reads otherwise antisense to a gene, 598 peaks and 5.8% of reads could not be related to annotated genes. Peaks were allowed up to 2000 bases downstream of the annotated 3’UTR end, but not inside another gene on the same strand. Statistically significant changes to gene expression, poly (A) tail-length and APA were calculated as previously described (Turner et al. 2020).

### Bioinformatic analyses

Bioinformatics for QKI HITS-CLIP was performed as previously described (Pillman et al. 2018). In brief, BAM files for samples that were prepared using the QKI-5 specific and panQKI antibodies were pooled together by strand. Peak calling was performed using the MACS2 with a cut off fold-enrichment value of 3 and a cut off q value of 0.05 (Zhang et al. 2008). Motif analysis was performed using HOMER (Heinz et al. 2010).

Publicly available QKI CLIP data (Hafner et al. 2010; Van Nostrand et al. 2016; Pillman et al. 2018) was used to generate the list of high confidence QKI-binding sites within 3’UTRs. The PolyA site atlas 2.0 (Herrmann et al. 2020) was used to identify productive cleavage and polyadenylation sites near QKI-binding sites.

Plots were generated using the Python packages Matplotlib and Seaborn, or GraphPad Prism. Motif logo figure was generated using the R package motifStack. Genome track figures were generated using Integrated Genomics Viewer (Thorvaldsdottir et al. 2013) or the University of Santa Cruz Genome Browser (Kent et al. 2002).

Gene ontology analysis was performed using the tool WEB-based GEne SeT AnaLysis Toolkit (WebGestalt) using Over-Representation Analysis with the databases Biological Process, Cellular Component and Molecular Function (Liao et al. 2019).

Expression of the long and short 3’UTR transcripts of COMT and TTF2, and QKI-5 mRNA were obtained from the Cancer Cell Line Encyclopedia breast cancer samples (Barretina et al. 2012). The identities of the long and short 3’UTR transcript were determined by inspection of the Ensembl genome browser, where ENST00000361682 was used as the long COMT transcript, ENST00000406520 as the short COMT transcript, ENST00000369466 as the long TTF2 transcript, ENST00000492682 as the short TTF2 transcript and ENST00000361752 as QKI-5. Scatterplots were generated using the Python package, Seaborn, and pearson correlation coefficients with associated P values were calculated using the Python package SciPy.

### Statistical analyses

A hypergeometric test was performed using the hypergeom.sf() function in the Python module scipy.stats to test for significant overlap between genes with QKI peaks and genes with APA in the PAT-Seq. This statistic takes into account the total number of expressed transcripts, which was determined to be 16,007 by counting the number of transcripts with a read count greater than 0 in the HMLE or mesHMLE RNA-Seq.

To test the changes in Distal PAS usage detected by qRT-PCR for statistical significance an ordinary 1-way ANOVA (p<0.05) was performed on all samples for each APA event followed by a Sidak’s multiple comparison post-hoc test using GraphPad PRISM software

## Supplementary Figure Legends

**Supplementary Figure 1: Validation of sequenced samples from PAT-Seq**

(A) A multidimensional scaling (MDS) plot depicting the clustering of the PAT-Seq samples.

(B) Venn diagrams showing overlap of APA events following EMT (HMLE-mesHMLE) or after QKI knockdown (mesHMLE-mesHMLE + QKI kd) (C) Categorical scatter plots showing gene expression data from the PAT-Seq as counts per million (CPM). Plots of the epithelial markers E-Cadherin and ESRP1, and the mesenchymal markers ZEB1, Vimentin and N-cadherin are shown to validate the induction of an EMT in HMLE after TGF-β treatment (mesHMLE). Plot of QKI expression is shown to validate changes in expression after EMT and after siRNA-mediated knockdown.

**Supplementary Figure 2: Overlap between CLIP-Seq replicates from same cell lines**

Venn diagrams showing overlap between (A) mesHMLE, (B) K562 and (C) HEPG2 CLIP-Seq experiments. HITS-CLIP-mesHMLE-rep-1 refers to a published experiment (Pillman et al. 2018) while HITS-CLIP-mesHMLE-rep-2 is the experiment performed in this study.

**Supplementary Figure 3: An alternative cleavage site downstream from the major cleavage site for QKI-5**

(A) UCSC Genome Browser image depicting expressed sequence tags (ESTs) within the QKI-5 3’UTR. Arrows indicate positions of the major upstream (QKI-5-Short) and alternate downstream (QKI-5-Long) cleavage sites. (B) Zoom in of the region of the QKI CLIP-seq peak at the major upstream cleavage site in the QKI-5 3’UTR. Boxes indicate the locations of the major QKI-5 polyadenylation site (PAS), and adjacent QKI sites.

**Supplementary Figure 4: Differential gene expression during EMT or following QKI knockdown**

(A,B) Volcano plots showing differential gene expression between HMLE-mesHMLE or mesHMLE-mesHMLE with QKI knockdown. Differentially expressed genes and gene numbers are highlighted in red. The dashed line represents an FDR cutoff of 0.05.

